# A method for genome editing in the anaerobic magnetotactic bacterium *Desulfovibrio magneticus* RS-1

**DOI:** 10.1101/375410

**Authors:** Carly R. Grant, Lilah Rahn-Lee, Kristen N. LeGault, Arash Komeili

## Abstract

**ABSTRACT:** Magnetosomes are complex bacterial organelles that serve as model systems for studying cell biology, biomineralization, and global iron cycling. Magnetosome biogenesis is primarily studied in two closely related Alphaproteobacterial *Magnetospirillum* spp. that form cubooctahedral-shaped magnetite crystals within a lipid membrane. However, chemically and structurally distinct magnetic particles have also been found in physiologically and phylogenetically diverse bacteria. Due to a lack of molecular genetic tools, the mechanistic diversity of magnetosome formation remains poorly understood. *Desulfovibrio magneticus* RS-1 is an anaerobic sulfate-reducing Deltaproteobacterium that forms bullet-shaped magnetite crystals. A recent forward genetic screen identified ten genes in the conserved magnetosome gene island of *D. magneticus* that are essential for its magnetic phenotype. However, this screen likely missed many interesting mutants with defects in crystal size, shape, and arrangement. Reverse genetics to target the remaining putative magnetosome genes using standard genetic methods of suicide vector integration has not been feasible due to low transconjugation efficiency. Here, we present a reverse genetic method for targeted mutagenesis in *D. magneticus* using a replicative plasmid. To test this method, we generated a mutant resistant to 5-fluorouracil by making a markerless deletion of the *upp* gene that encodes uracil phosphoribosyltransferase. We also used this method for targeted marker exchange mutagenesis by replacing *kupM,* a gene identified in our previous screen as a magnetosome formation factor, with a streptomycin resistance cassette. Overall, our results show that targeted mutagenesis using a replicative plasmid is effective in *D. magneticus* and may also be applied to other genetically recalcitrant bacteria.

**IMPORTANCE:** Magnetotactic bacteria (MTB) are a group of organisms that form small, intracellular magnetic crystals though a complex process involving lipid and protein scaffolds. These magnetic crystals and their lipid membrane, termed magnetosomes, are model systems for studying bacterial cell biology and biomineralization as well as potential platforms for biotechnological applications. Due to a lack of genetic tools and unculturable representatives, the mechanisms of magnetosome formation in phylogenetically deeply-branching MTB remain unknown. These MTB contain elongated bullet-/tooth-shaped magnetite and greigite crystals that likely form in a manner distinct from the cubooctahedral-shaped magnetite crystals of the genetically tractable Alphaproteobacteria MTB. Here, we present a method for genome editing in the anaerobic Deltaproteobacterium *Desulfovibrio magneticus* RS-1, the first cultured representative of the deeply-branching MTB. This marks a crucial step in developing *D. magneticus* as a model for studying diverse mechanisms of magnetic particle formation by MTB.

## INTRODUCTION

Magnetotactic bacteria (MTB) are a group of diverse microorganisms that align along magnetic fields via their intracellular chains of magnetic crystals (1, 2). Each magnetic crystal consists of either magnetite (Fe_3_O_4_) or greigite (Fe_3_S_4_) and is synthesized within a complex organelle called a magnetosome (3). The first cultured MTB were microaerophilic Alphaproteobacteria, which form cubooctahedral-shaped magnetite crystals, and have served as model organisms for understanding magnetosome formation (4–7). Early studies on *Magnetospirillum* spp. found a lipid-bilayer membrane, with a unique suite of proteins, surrounding each magnetite crystal (8–10). Development of genetic tools in *Magnetospirillum magneticum* AMB-1 and *Magnetospirillum gryphiswaldense* MSR-1 revealed a conserved *<u>ma</u>*gnetosome gene island (MAI) that contains the factors necessary and sufficient for the formation of the magnetosome membrane, magnetite biomineralization within the lumen of the magnetosome, and alignment of the magnetosomes in a chain along the length of the cell (3, 11). These molecular advances, along with the magnetic properties of magnetosomes, have made MTB ideal models for the study of compartmentalization and biomineralization in bacteria as well as a target for the development of biomedical and industrial applications.

Improvements in isolation techniques and sequencing have revealed that MTB are ubiquitous in many aquatic environments. Based on phylogeny and magnetosome morphology, MTB can be categorized into two subgroups. The first subgroup includes members of the Proteobacteria with similar magnetosome morphology to *Magnetospirillum* spp. The second subgroup comprises MTB from more deep-branching lineages, including members of the Deltaproteobacteria, Nitrospirae, and Omnitrophica phyla, which synthesize elongated bullet-/tooth-shaped magnetite and/or greigite crystals (12, 13). While all MTB sequenced to date have their putative magnetosome genes arranged in a distinct region of their genome (3, 14–16), many of the genes essential for magnetosome biogenesis in *Magnetospirillum* spp. are missing from the genomes of deep-branching MTB (13). Likewise, a conserved set of *mad* (*<u>m</u>*agnetosome *<u>a</u>*ssociated *<u>D</u>*eltaproteobacteria) genes are only found in deep-branching MTB (13, 17–19). This suggests a genetic diversity underpinning the control of magnetosome morphology and physiology in non-model MTB that is distinct from the well-characterized *Magnetospirillum* spp.

*Desulfovibrio magneticus* RS-1, the first cultured MTB outside of the Alphaproteobacteria, is an anaerobic sulfate-reducing Deltaproteobacterium that forms irregular bullet-shaped crystals of magnetite (20, 21). As with the *Magnetospirillum* spp., the magnetosome genes of *D. magneticus* are located within a MAI and include homologs to some *mam* genes as well as *mad* genes (13, 17, 22). Recently, we used a forward genetic screen combining random chemical and UV mutagenesis with whole genome resequencing to identify mutations in ten *mam* and *mad* genes of the *D. magneticus* MAI that resulted in non-magnetic phenotypes (19). However, this screen relied on a strict selection scheme for nonmagnetic mutants. As such, we likely missed magnetosome genes that are important for regulating the shape, size, and arrangement of magnetosomes. In order to elucidate the degree of conservation between *mam* genes and determine a function for the *mad* genes in *D. magneticus*, a reverse genetic method for targeted mutagenesis is necessary.

*Desulfovibrio* spp. have gained much attention for their important roles in the global cycling of numerous elements, in biocorrosion, and in the bioremediation of toxic metal ions (23, 24). Development of genetic tools, such as expression vectors, transposons, and targeted genome editing systems, has allowed for a more detailed examination of the important activities of a few *Desulfovibrio* spp. (25, 26). Targeted mutagenesis using a one-step double recombination method was first achieved in *Desulfovibrio fructosovorans* and, more recently, in *Desulfovibrio gigas* and *Desulfovibrio desulfuricans* ND132 (27–29). In this method, plasmids that are electroporated into the cell are thought to be rapidly linearized by endogenous restriction-modification systems (29–31). The linearized plasmid DNA, carrying a selectable marker flanked by upstream and downstream regions of homology to a target gene, can then undergo double recombination into the chromosome in one step (**Fig. 1A**). This efficient one-step method, which is dependent on electroporation of the plasmid (27–29), is unlikely to be applicable in *D. magneticus* because plasmid uptake has only been demonstrated using conjugal transfer (19). The second targeted mutagenesis method, used in *Desulfovibrio vulgaris* Hildenborough, is a two-step double recombination that makes use of a non-replicative, or suicide, vector (30, 31). In the first step of this method, a suicide vector, with sequences upstream and downstream of the target gene, integrates into the genome upon the first homologous recombination event (**Fig. 1B**). Next, a second recombination event occurs whereby the vector is excised from the genome and cells with the desired genotype are selected with an antibiotic marker and/or a counterselection marker (30, 31) (**Fig. 1B**). For many bacteria, including *D. magneticus,* plasmid uptake and integration occur at low enough frequencies to prevent the use of suicide vectors for genetic manipulation (19).

**Figure 1.**
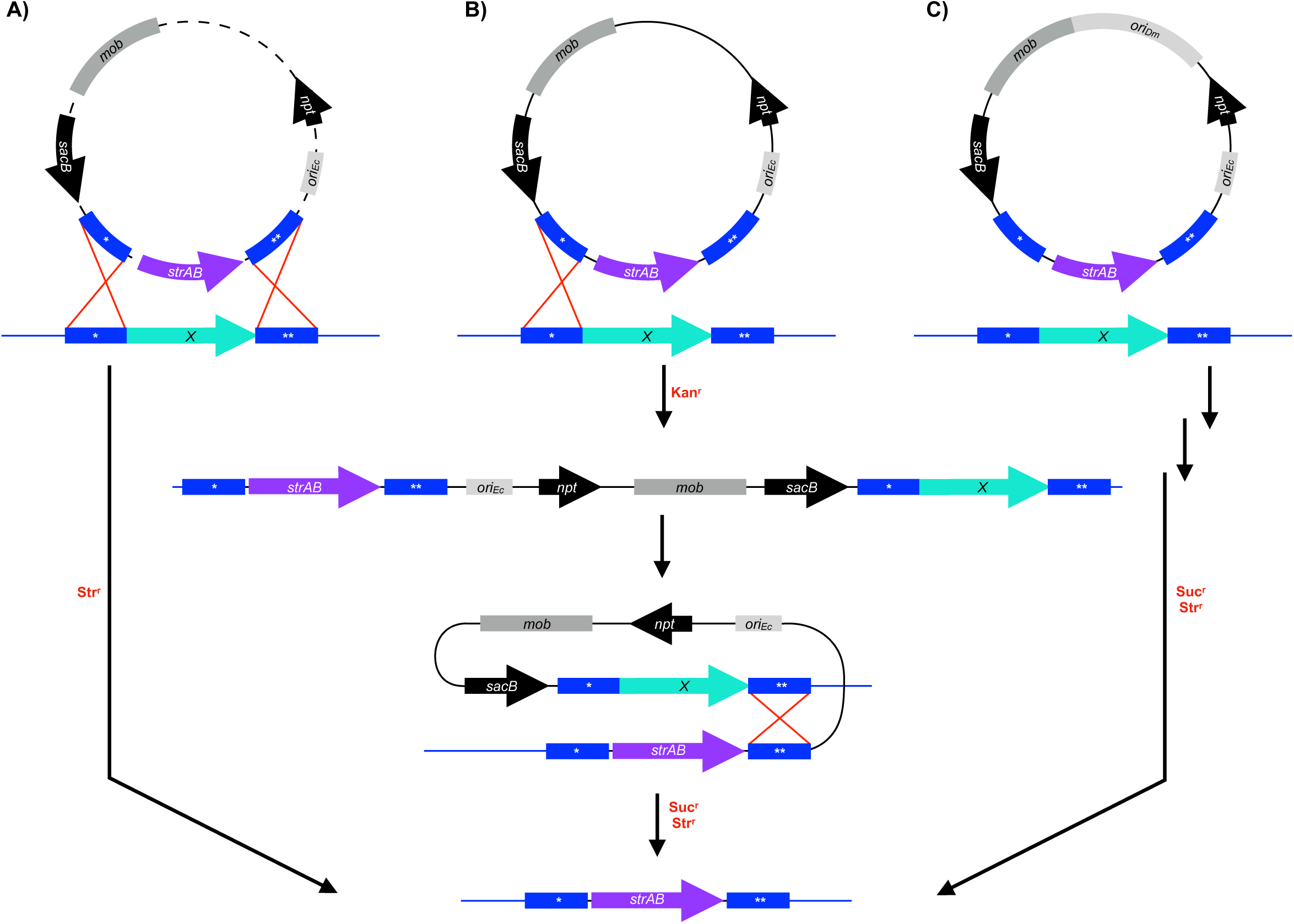
Schematic of deletion methods used in *Desulfovibrio* spp. (**A-C**) Plasmids (dashed lines) are designed to replace a target gene (*X*, aqua arrow) in the chromosome (blue line) with a streptomycin resistance cassette (*strAB*, purple arrow). Regions upstream (*) and downstream (**) of the target gene (blue boxes) on the chromosome undergo recombination (red lines) with homologous regions that are cloned into the deletion plasmid. Key steps, such as recombination events (red crosses), are indicated in the boxes and the selection steps are labeled in red. (**A**) Double recombination can occur in one step after plasmids are linearized by endogenous restriction enzymes. Mutants are selected using the marker (*e.g. strAB*) that was exchanged with the target gene. (**B**) Two-step double recombination is possible when suicide vectors integrate into the chromosome in the first homologous recombination event and then recombine out after the second homologous recombination event. The first step and second step are selected for with antibiotic resistance markers (*e.g. npt*) and counterselectable markers (*e.g. sacB),* respectively. (**C**) A replicative deletion plasmid designed to target genes for deletion may undergo double recombination in one or two steps as shown in A and B, respectively. After passaging the cells without antibiotic, mutants are selected with an antibiotic resistance cassette (*e.g. strAB*) and a counterselectable marker (*e.g. sacB). mob,* mobilization genes (*mobA\ mobB, mobC*) and *oriT; npt,* kanamycin-resistance gene; *ori_Dm_,* origin of replication for *D. magneticus; ori_Ec_,* origin of replication for *E. coli.*

Here, we develop a method for targeted gene deletion using a replicative plasmid, thereby bypassing the need for suicide vector integration (**Fig. 1C**). We generated a mutant resistant to 5-fluorouracil by making a markerless deletion of the *upp* gene which encodes an enzyme in the pyrimidine salvage pathway that is nonessential under standard laboratory conditions. Additionally, we deleted *kupM,* a gene encoding a potassium transporter that acts as a magnetosome formation factor (19), via marker exchange with a streptomycin resistance cassette. Deletion of both *upp* and *kupM* conferred the expected phenotypes, which were subsequently complemented *in trans.* Overall, our results show that targeted mutagenesis using a replicative plasmid is possible in *D. magneticus.* It may also be suitable for other bacteria for which replicative plasmid uptake is possible, but at a rate too low for suicide vector integration.

## RESULTS

### Design of a replicative deletion plasmid using *sacB* counterselection

Targeted genetic manipulation in most bacteria requires a method to efficiently deliver foreign DNA destined for integration into the chromosome. One commonly used method involves suicide vector uptake and integration prior to the first selection step (**Fig. 1B**). In *D. magneticus*, plasmid transfer has only been achieved via conjugation at low efficiencies making the uptake and subsequent integration of suicide vectors into its chromosome an unlikely event (19). As such, we hypothesized that we could bypass the need for a suicide vector by using a stable, replicative plasmid designed to delete specific genes via homologous recombination (**Fig. 1C**). Two features of this method will allow for isolation of desired mutants: (1) a selectable marker is used to identify double recombination events at the targeted site and (2) a counterselectable marker distinguishes the desired mutant cells, which have lost all remaining copies of the plasmid.

*sacB* is a common counterselection marker that is effective in many bacteria. The *sacB* gene from *Bacillus subtilis* encodes levansucrase, which converts sucrose to levans that are lethal to many Gram-negative bacteria, including *D. vulgaris* Hildenborough (30, 32, 33). To test its functionality in *D. magneticus,* we inserted *sacB* under the expression of the *mamA* promoter in a plasmid that replicates in both *Escherichia coli* and *D. magneticus* (**Fig. 2A**). This plasmid (pAK914) and a control plasmid were then conjugated into *D. magneticus*. We found no growth inhibition for *D. magneticus* cells with the control plasmid in the presence of sucrose and kanamycin. In contrast, cells expressing *sacB* were unable to grow with kanamycin and sucrose concentrations of 1% (w/v) or higher (data not shown). To test if plasmids could be cured, *D. magneticus* with pAK914 was passaged two times in liquid media containing no antibiotic and plated on 1% sucrose. Individual sucrose resistant (Suc^r^) colonies were inoculated and screened for kanamycin sensitivity (Kan^s^). All isolated colonies (n=16) were Kan^s^, suggesting that the cells had lost the plasmid. These experiments demonstrate that *sacB* is a suitable counterselection marker in *D. magneticus*.

**Figure 2.**
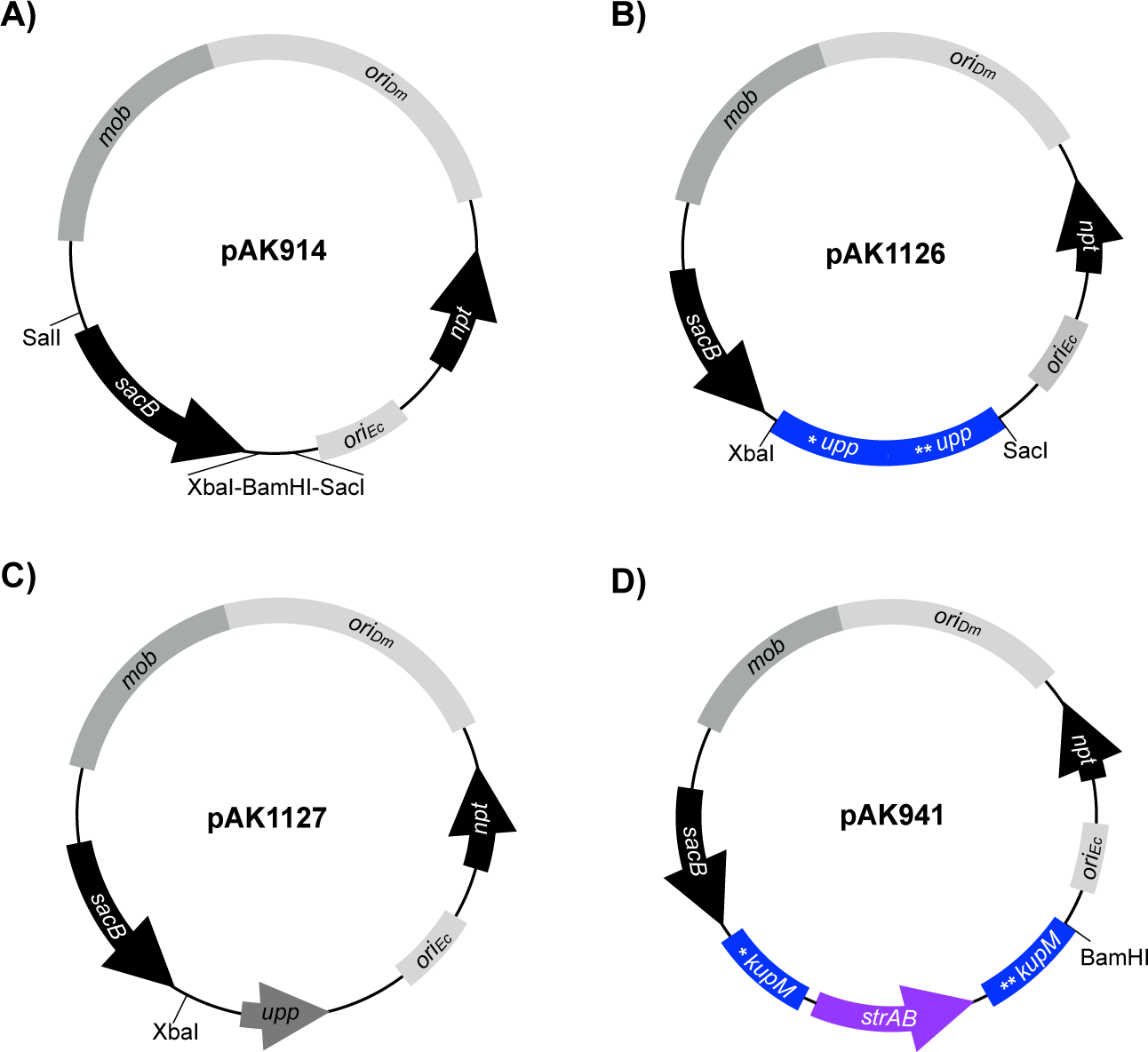
Plasmids constructed for the present study. (**A**) Expression plasmid pAK914 expresses *sacB* from the *mamA* promoter and is the parent vector for the deletion plasmids and *upp* expression plasmid described below. (**B**) Replicative deletion plasmid to target *upp* for markerless deletion. The *upp* deletion cassette was cloned into XbaI-SacI of pAK914. (**C**) Expression plasmid used for *upp* complementation. The *upp* gene and its promoter were cloned into BamHI-SacI of pAK914. (**D**) Replicative deletion plasmid to target *kupM* for marker exchange mutagenesis with *strAB.* The *kupM::strAB* deletion cassette was cloned into XbaI of pAK914. Labeling and colors correspond to **Figure 1**.

### Construction of a Δupp strain by markerless deletion

To test our replicative deletion method, we chose to target the *upp* gene, the mutation of which has a selectable phenotype. The *upp* gene encodes uracil phosphoribosyltransferase (UPRTase), a key enzyme in the pyrimidine salvage pathway that catalyzes the reaction of uracil with 5-phosphoribosyl-α-1-pyrophosphate (PRPP) to UMP and PP/ (34) (**Fig. 3A**). When given the pyrimidine analog 5-fluorouracil (5-FU), UPRTase catalyzes the production of 5-fluoroxyuridine monophosphate (5-FUMP). 5-FUMP is further metablized and incorporated into DNA, RNA, and sugar nucleotides resulting in eventual cell death (**Fig. 3A**) (35, 36). Previous studies have shown that *Δupp* mutants of *D. vulgaris* Hildenborough are resistant to 5-FU while wild type (WT) cells are effectively killed by the pyrimidine analog (31, 37). The *D. magneticus* genome has a homolog (*DMR_08390*) to the *D. vulgaris* Hildenborough *upp* gene that is likely functional as detected by the sensitivity of *D. magneticus* to 5-FU (**Fig. 3B, Fig. 4A**). To show that the *upp* gene product confers 5-FU sensitivity and to validate our replicative deletion system, we chose to target the *D. magneticus upp* gene for markerless deletion.

**Figure 3.**
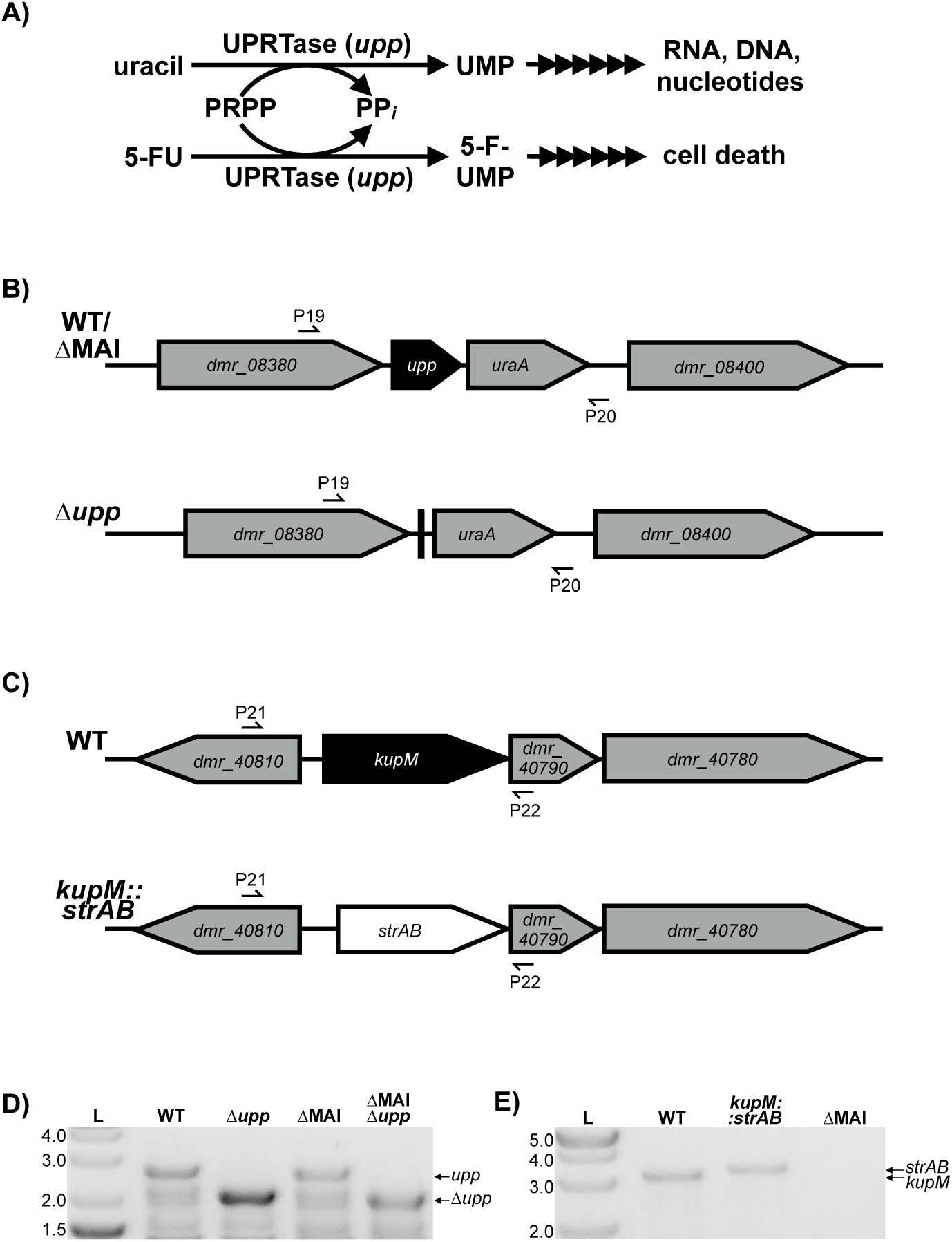
(**A**) The *upp* gene encodes UPRTase which is a key enzyme in the uracil salvage pathway. The product of the UPRTase reaction, UMP, is processed by downstream enzymes in pathways for RNA, DNA, and sugar nucleotide synthesis. 5-FU causes cell death by incorporating into this pathway via UPRTase. (**B**) Schematic of genomic region of *upp* in WT or ΔMAI (top) and the △upp mutant (bottom). (**C**) Genomic region of *kup* in WT (top) and *kup::strAB* (bottom). Primers used to screen for the correct genotype are indicated with half arrows. (**D**) *Δupp* mutants in WT and ΔMAI backgrounds were confirmed by PCR using primers P19/P20 and agarose gel electrophoresis. WT and ΔMAI show a band corresponding to the *upp* gene (2691 bp) while the *Δupp* mutants have a smaller band corresponding to a markerless deletion of the *upp* gene (2079 bp). The lower bands are likely non-specific PCR products. (**E**) *kupM::strAB* genotype confirmation by PCR and agarose gel electrophoresis using primers P21/P22 (WT=3069 bp; *kupM::strAB=3263* bp, ΔMAI=N/A).

**Figure 4.**
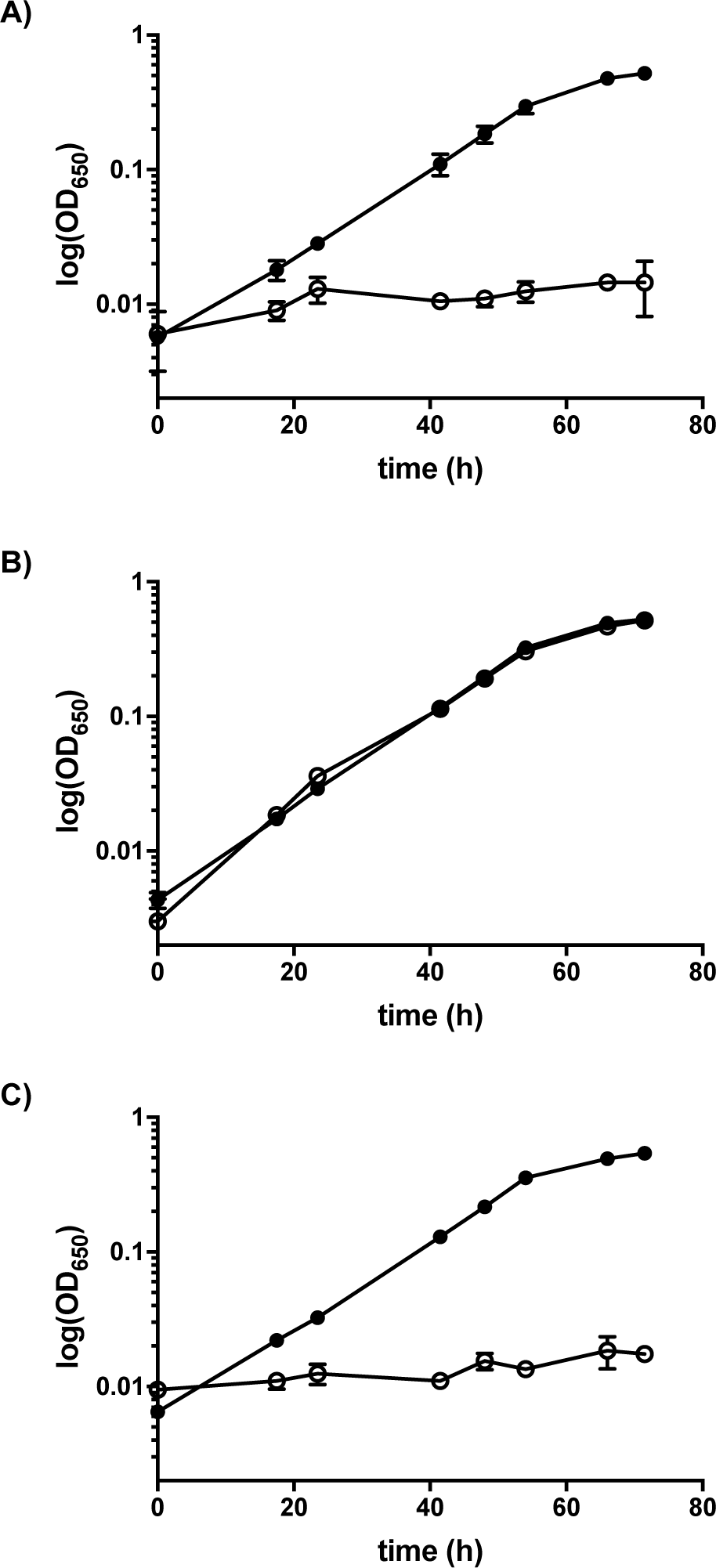
*upp* mutant and complementation phenotype. Growth of the parent strain (△MAI) (**A**), *upp* deletion (△MAI △upp) (**B**), and complementation of the *upp* deletion (△MAI *△upp/upp+*) (**C**) when grown with 1.25 μg/ml 5-FU (squares) or without 5-FU (circles). Data presented are averages from 2-3 independent cultures; error bars indicate the standard deviation.

To construct a *upp* deletion vector, a markerless cassette containing the regions upstream and downstream of the *upp* gene were inserted into plasmid pAK914 (**Fig. 2B**). The resulting plasmid (pAK1126) was transferred to WT *D. magneticus* and a non-magnetic strain (ΔMAI) (19) by conjugation and single, kanamycin resistant (Kan^r^) colonies were isolated and passaged in growth medium containing no antibiotic. After the third passage, *upp* mutants that had lost the vector backbone were selected for with 5-FU and sucrose. Compared with a control plasmid (pAK914), over 10-fold more 5-FU resistant (5-FU^r^) mutants were obtained from pAK1126. PCR of the region flanking the *upp* gene confirmed that the 5-FU^r^ colonies from pAK1126 resulted in markerless deletion of *upp* (Δupp) while 5-FU^r^ colonies from pAK914 were likely the result of point mutations (**Fig. 3B, Fig. 3D**). Similar to results obtained for *D. vulgaris* Hildenborough (31), the *Δupp* mutant of *D. magneticus* was able to grow in the presence of 5-FU (**Fig. 4B, Table 2**). Complementation of the *upp* gene *in trans* restored UPRTase function and cells were no longer able to grow with 5-FU (**Fig. 2C, Fig. 4C, Table 2**). These experiments demonstrate that a replicative plasmid can be used to directly edit the *D. magneticus* genome.

### Construction of a *Δkup* strain by marker exchange mutagenesis

Because many genetic mutations confer no selectable phenotype, we sought to develop our replicative deletion plasmid for marker exchange mutagenesis. To test this system, we chose to replace a gene with a known phenotype, *kupM (DMR_40800),* with a streptomycin-resistance gene cassette (*strAB). kupM* is located in the *D. magneticus* MAI and encodes a functional potassium transporter (19). Mutant alleles in *kupM,* including missense, nonsense, and frameshift mutations, were previously identified in our screen for non-magnetic mutants (19). These *kupM* mutations resulted in cells that rarely contained electron-dense particles and were unable to turn in a magnetic field, as measured by the coefficient of magnetism or C_mag_ (19).

To mutate *kupM,* we inserted a marker-exchange cassette, with regions upstream and downstream of *kupM* flanking *strAB,* into pAK914 (**Fig. 2D**) to create the deletion plasmid pAK941. Following conjugation, single colonies of *D. magneticus* with pAK941 were isolated by kanamycin selection. After three passages in growth medium without selection, potential mutants were isolated on plates containing streptomycin and sucrose. Single colonies (n=48) that were streptomycin resistant (Str^r^) and Suc^r^ were inoculated into liquid medium and screened for Kan^s^. Of the ten isolates that were Kan^s^, two had the correct genotype (*ΔkupM::strAB*) as confirmed by PCR and sequencing (**Fig. 3C, Fig. 3E**).

Similar to the phenotypes previously observed in *kupM* mutants (19), *ΔkupM::strAB* cells were severely defective in magnetosome synthesis and ability to turn in a magnetic field (**Fig. 5**). Though a slight C_mag_ could be measured, few cells contained electron-dense particles or magnetosomes. Importantly, the WT phenotype was rescued by expressing *kupM* on a plasmid in the *ΔkupM::strAB* mutant (**Fig. 5**). These results confirm that the replicative deletion plasmid method described here can be used successfully for marker exchange mutagenesis.

**Figure 5.**
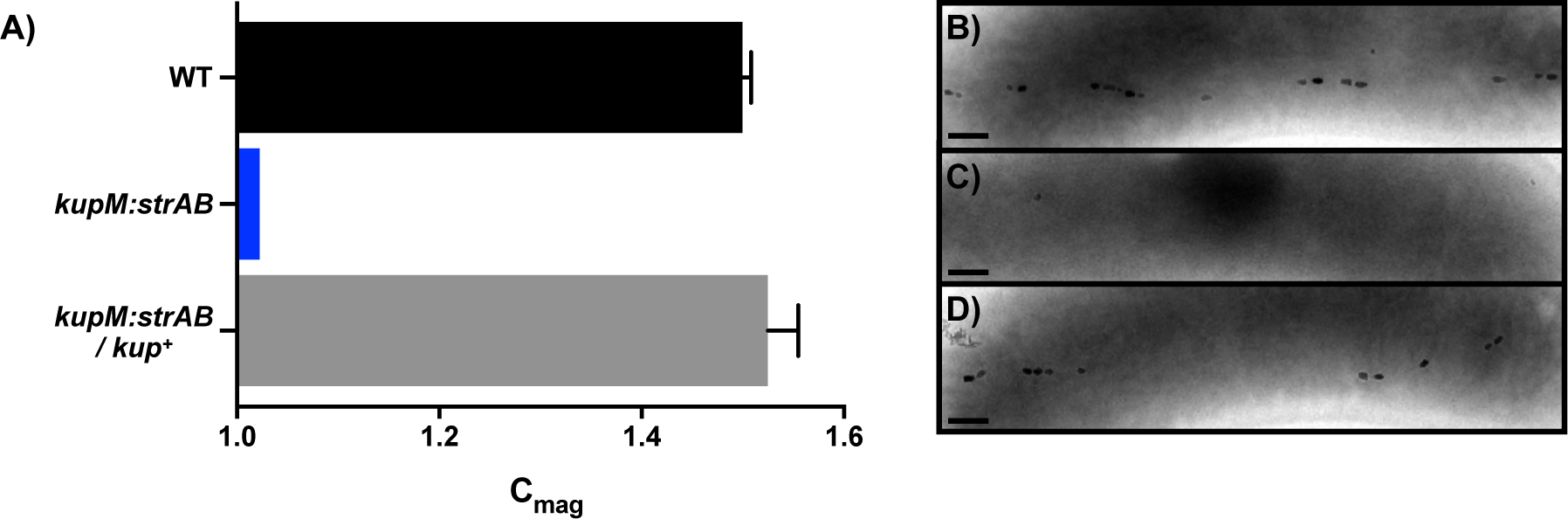
*kupM* mutant and complementation phenotype. C_mag_ values (**A**) and electron micrographs of WT (**B**), *kupM::strAB* (**C**), and *△kupM::strAB/kup+* (**D**). Scale bars, 200 nm. Data presented are averages from 4 independent cultures; error bars indicate the standard deviation.

## DISCUSSION

In this study, we expand the genetic toolbox of *D. magneticus* to include a replicative plasmid method for targeted mutagenesis (**Fig. 1C**). We show the utility of this method for markerless deletion of genes with a selectable phenotype and for marker exchange mutagenesis. Some of the earliest examples of targeted mutagenesis in Gram-negative bacteria used replicative plasmids, similar to the method described here. (33, 38). These studies, which predated the application of suicide vectors, relied on plasmid instability by introducing a second plasmid of the same incompatibility group or by limiting nutrients in the growth medium (33, 38).

Because the *D. magneticus* genetic toolbox has a limited number of plasmids, antibiotic markers, and narrow growth constraints, we used a replicative plasmid and established *sacB* as a counterselection marker to generate and isolate mutants. While *sacB* counterselection was ultimately successful, a large number of false-positives were also isolated at the sucrose selection step. Mutations in *sacB* have been found to occur at a high frequency in many bacteria (30, 39–42). Indeed, we found that deletions and mutations in *P_mamA__sacB* are abundant in the false-positive Suc^r^ Str^r^ isolates (data not shown). Alternative counterselection markers, including *upp,* have been shown to select for fewer false-positives (31, 42–44). Since *D. magneticus* is sensitive to 5-FU only when the *upp* gene is present (**Fig. 4**), the *upp* mutants generated in this study may be used as the parent strains for future targeted mutagenesis using *upp* as a counterselectable marker rather than *sacB.* Additionally, the combined use of *upp* and *sacB* for counterselection could reduce the false-positive background that results from the accumulation of mutations in these markers.

The replicative deletion plasmid described here is designed to replace a target gene with an antibiotic resistance marker. As such, the construction of strains with multiple directed mutations will be complicated by the need for additional antibiotic-resistance markers, which are limited in *D. magneticus.* These limitations may be overcome by removing the chromosomal antibiotic marker in subsequent steps (33, 45, 46). Ultimately, improvements in conjugation efficiency or methods for electroporation with high transformation efficiency are desired. Similar to the ongoing development of genetics in *D. vulgaris* Hildenborough, establishment of a suicide vector delivery system in *D. magneticus* will allow for more high-throughput targeted mutagenesis and even the construction of markerless deletion mutants (25, 31).

Overall, we have demonstrated the utility of a replicative deletion plasmid used to generate the first targeted mutants of *D. magneticus*. This method marks a crucial step in developing *D. magneticus* as a model for the study of anaerobic sulfate reduction and diverse mechanisms of magnetic particle formation by MTB. Both MTB and sulfate-reducing bacteria have been singled out for their role in the global cycling of numerous elements and for potential applications, such as bioremediation (23, 24, 47, 48). *D. magneticus*, in particular, may be useful in the bioremediation of heavy-metals and in the global cycling of iron, since it can form both magnetosomes and other iron-containing organelles (49, 50). Through genetic manipulation of *D. magneticus*, pathways of elemental cycling and heavy-metal turnover may now be explored. Additionally, genetic manipulation of *D. magneticus* will further our understanding of magnetosome formation and provide answers to many longstanding questions for the deeply-branching MTB: Which proteins regulate and control magnetosome formation? To what extent are lipid membranes involved in forming these crystals? How is the elongated and irregular crystal shape achieved? Finally, in addition to *D. magneticus*, the method described here may extend to other bacteria that are not amenable to targeted mutagenesis with suicide vectors but are able to accommodate replicative plasmids.

## MATERIALS AND METHODS

### Strains, media, and growth conditions

The bacterial strains used in this study are listed in **Table 1**. All *E. coli* strains were cultured aerobically with continuous shaking at 250 RPM at 37°C in lysogeny broth (LB). *D. magneticus* strains were grown anaerobically at 30°C in sealed Balch tubes with a N_2_ headspace containing RS-1 Growth Medium (RGM) that was degassed with N_2_, unless otherwise stated (50). Sodium pyruvate (10 mM) was used as an electron donor with fumaric acid disodium (10 mM) as the terminal electron acceptor. RGM was buffered with Hepes and the pH was adjusted to 6.7 with NaOH (19). Before inoculating with cells, RGM was supplemented with 0.8% (v/v) Wolfe’s vitamins, 100 μM ferric malate, and 285 μM cysteine-HCl (50). Solid agar plates were prepared by adding 1.5% agar (wt/vol) to LB and 1% agar (wt/vol) to RGM. Vitamins (0.8% v/v), ferric malate (20 μM), and cysteine (285 μM) were added to the molten RGM agar, as well as antibiotics and selective agents, as needed. For *D. magneticus,* all plating steps were carried out aerobically, transferred to an anaerobic jar, and incubated at 30°C for 10-14 days, as described previously (19). Antibiotics and selective agents used are as follows: kanamycin (50 μg/mL for *E. coli* strains, 125 μg/ml for *D. magneticus* strains), streptomycin (50 μg/ml for *E. coli* and *D. magneticus* strains), diaminopimelic acid (300 μM for *E. coli* WM3064), 5-FU (2.5 μg/ml for *D. magneticus* strains), and sucrose (1% for *D. magneticus* strains).

**TABLE 1.**
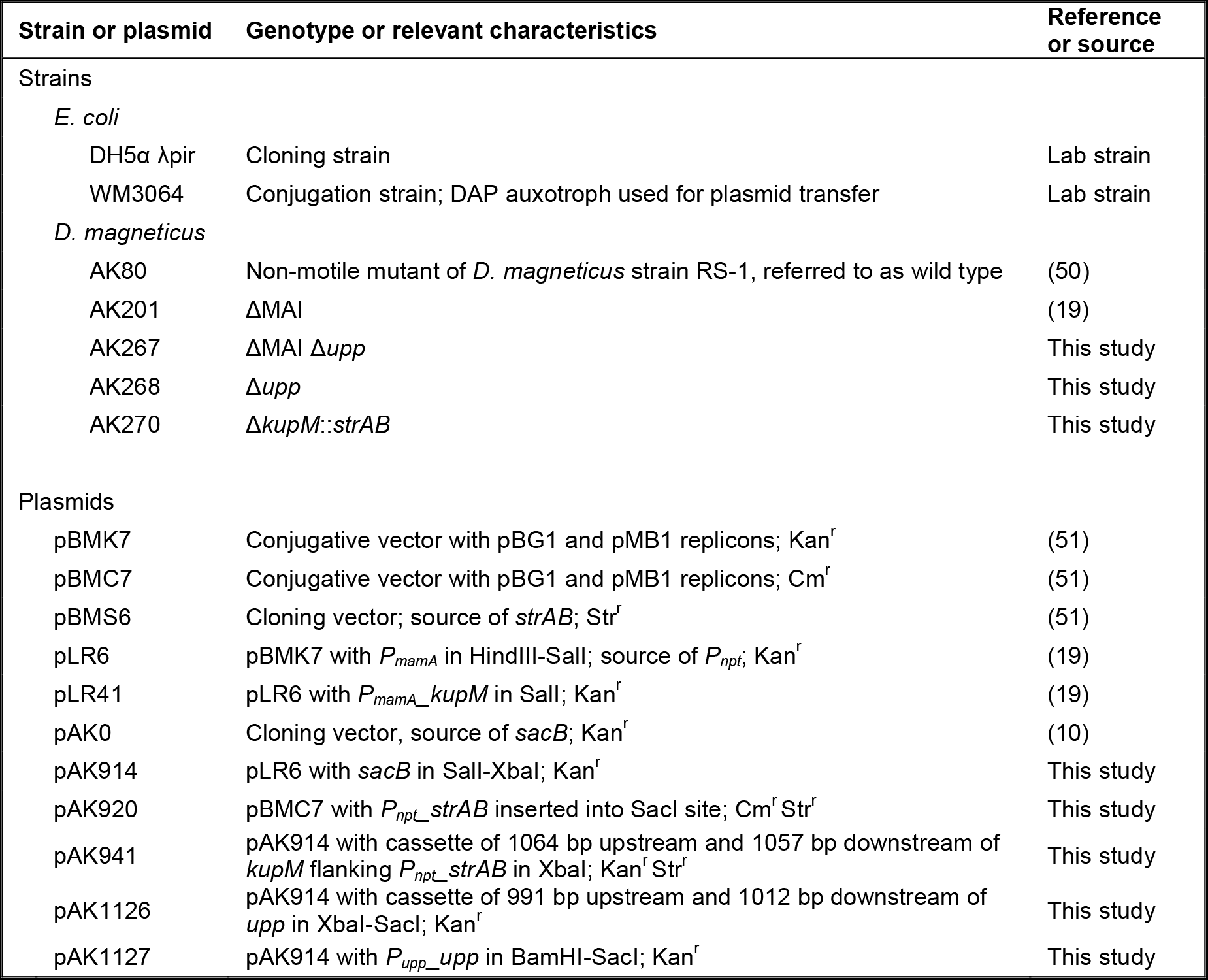
Bacterial strains and plasmids used in this study.

### Plasmids and cloning

All plasmids used in this work are listed in **Table 1**. All cloning was performed in *E. coli* DH5α λpir using the Gibson method or restriction enzyme ligation (Gibson, et al 2009). For PCR amplification, KOD (EMD Millipore, Germany) and GoTaq (Promega, USA) DNA polymerases were used with the primers listed in Table S1. All upstream and downstream homology regions were amplified from *D. magneticus* genomic DNA. *strAB* and *P_npt_* were amplified from pBMS6 and pLR6, respectively, and subcloned into pBMC7 to make pAK920 which served as the template for amplifying *Pnpt_strAB* for the deletion vectors. *sacB* was amplified from pAK0 and inserted into pLR6 digested with SalI and XbaI to create pAK914. To construct a plasmid for the targeted deletion of *upp (DMR_08390),* 991 bp upstream and 1012 bp downstream of *upp* were amplified and inserted into pAK914 digested with XbaI and SacI using a 3-piece Gibson assembly. To create the *upp* complementation plasmid, pAK914 was digested with BamHI and SacI and the *upp* gene, with its promoter, were PCR amplified from *D. magneticus* genomic DNA. To construct pAK941 for marker exchange mutagenesis of *kupM*, a cassette of 1064 bp upstream region and 1057 bp downstream region flanking *P_npt__strAB* was assembled using Gibson cloning. The cassette was amplified and inserted into pAK914 digested with XbaI using a two-piece Gibson assembly.

### *upp* and *kup* mutant generation and complementation

Replicative deletion plasmids were transformed into *E. coli* WM3064 by heat shock and transferred to *D. magneticus* by conjugation, as described previously (19). Single colonies of Kan^r^ *D. magneticus* were isolated and inoculated into RGM containing no antibiotic. Cultures were passaged three times and spread on 1% agar RGM plates containing either 50 μg/ml streptomycin and 1% sucrose or 2.5 μg/ml 5-FU and 1% sucrose. Single colonies were screened for Kan^s^ and by PCR using the primers listed in **Table 2**. Successful *upp* and *kup* mutants were confirmed by Sanger sequencing. Expression plasmids for the complementation of *Δkup::strAB* and *Δupp* as well as empty vectors for controls were transferred to *D. magneticus* strains as described above. Transconjugants were inoculated into RGM containing kanamycin in order to maintain the plasmids.

**TABLE 2.** Growth rates and generation times of the parent strain (△MAI), *△upp* mutant, and *upp* complementation in *trans* with and without treatment with 5-FU.

| Strain | growth rate (h<sup>-1</sup>) |  | generation time (h) |  |
| --- | --- | --- | --- | --- |
|  | - 5-FU | + 5-FU | - 5-FU | + 5-FU |
| ΔMAI | 0.077 ± 0.0017 | N/A | 9.1 ± 0.2 | N/A |
| ΔMAI Δupp | 0.079 ± 0.0017 | 0.070 ± 0.0040 | 8.8 ± 0.2 | 10.0 ± 0.6 |
| ΔMAI Δupp/upp<sup>+</sup> | 0.076 ± 0.0041 | N/A | 9.1 ± 0.5 | N/A |

### Mutant phenotype and complementation analyses

The growth and coefficient of magnetism (C_mag_) of *D. magneticus* strains were measured in a Spec20 spectrophotometer at an optical density of 650 nm (OD_650_), as described previously (10, 50). For *upp* mutant and complementation analysis, RGM was supplemented with 5-FU (2.5 μg/ml in 0.1% DMSO) or DMSO (0.1%) and growth was measured for WT and *Δupp* strains with an empty vector (pAK914) and for the *Δupp* strain with the complementation plasmid pAK1127. For *kup* mutant and complementation analysis, the Cmag was measured by placing a large bar magnet parallel or perpendicular to the sample in order to measure the maximum or minimum absorbance, respectively, as the *D. magneticus* strains rotate 90 degrees with the magnetic field. The ratio of maximum to minimum absorbances was calculated as the C_mag_ (10). Whole-cell transmission electron microscopy (TEM) was performed as previously described (50). The Cmag calculations and TEM were performed for WT *D. magneticus* with an empty vector (pBMK7) and *Δkup::strAB* with an empty vector (pBMK7) or complementation plasmid (pLR41). For all growth measurements, C_mag_ measurements, and TEM, plasmids were maintained in cells with 125 μg/ml kanamycin.

## ACKNOWLEDGMENTS

We would like to thank the members of the Komeili lab for helpful suggestions and discussions and the staff at the University of California Berkeley Electron Microscope Laboratory for advice and assistance in electron microscopy sample preparation and data collection.

